# MAPK activity and MAP kinase phosphatase 2 (AtMKP2) protein stability under altered glutathione homeostasis in *Arabidopsis thaliana*

**DOI:** 10.1101/2024.01.15.575793

**Authors:** Yingxue Yang, Sanja Matern, Heike Steininger, Marcos Hamborg Vinde, Thomas Rausch, Tanja Peskan-Berghöfer

## Abstract

Mitogen-activated protein kinases (MAPKs) are important signaling players involved in various responses to diverse environmental stresses. MAP kinase phosphatases (MKPs) are crucial negative regulators of MAPKs and control the intensity and duration of MAPK activation. It has been shown that transgenic tobacco plants with increased glutathione content display an oxidative shift and have constitutively active immunity-related MAPKs. The mechanism by which glutathione can activate or keep these MAPKs in activated state is unclear. In this study, it is shown that the *Arabidopsis* stress-related MAPKs, AtMPK3 and AtMPK6 are hypersensitive to a pathogen-associated molecular pattern flg22 in the *cat2-1* line, under the conditions causing an altered glutathione homeostasis and elevated oxidative stress responses in this background. As AtMKP2 is the only dual specificity phosphatase deactivating AtMPK3 and AtMPK6 in response to oxidative stress, the stability of the wild-type AtMKP2 protein and the mutant version of the protein with the substitution of the cys109 in the active site with serine has been studied in wild type (Col-0) and *cat2-1* background. The results indicate that AtMKP2 is a stable protein in both genetic backgrounds, whereas the active site cys109 stabilizes the protein under severe oxidative stress conditions and can be glutathionylated *in vitro*.

## Introduction

As sessile organisms, plants are continuously exposed to different types of environmental challenges, which impact plant growth and yield. The signals triggered by environmental stimuli are transmitted *via* highly coordinated signaling system, initiating thereby adaptive responses which promote plant survival. MAPK cascades are pivotal players in this signaling network (Zhang and Zhang, 2022). They are composed of at least three protein-kinases in a row including MAP kinase kinase kinase (MAPKKK), MAP kinase kinase (MAPKK) and MAP kinase (MAPK), which in subsequent phosphorylation events activate each other, conducting the signal to the downstream targets (Zhang et al., 2018). MAP kinases (MAPKs) represent the terminal components of the cascade, which are activated by dual phosphorylation on conserved threonine and tyrosine residues in the TxY motif of the activation loop (Jonak et al., 2002). These kinases function as molecular switches that turn on the expression of specific sets of genes, resulting in the activation of cellular responses (Ma and Nicolet, 2023).

The outcome of the signaling mediated by the MAPK cascades depends on signal magnitude and duration, which rely on regulated deactivation of MAPKs by phosphatases (Bartels et al., 2010; Lin et al., 2022). Both, the serine/threonine phosphatases and protein tyrosine phosphatases (PTPs) can regulate plant MAPKs by dephosphorylating either threonine or tyrosine residues in the active site. However, only a particular sub-group of PTPs, the dual-specificity phosphatases (DSPs), can dephosphorylate both tyrosine and threonine residues within the activation loop (Jiang et al., 2018). Five out of twenty-two DSPs identified in *Arabidopsis* genome contain the active site sequence motif characteristic for mammalian MAPK phosphatases and have been confirmed to interact and dephosphorylate plant MAPKs (Bartels et al., 2010; Jiang et al., 2018). However, only two of them, MAPK phosphatases 1 and 2 (AtMKP1 and AtMKP2) are able to interact with and deactivate the MAPKs AtMKP3 and AtMKP6, which play a pivotal role in plant responses to pathogens and abiotic stresses (Anderson et al., 2011; Besteiro and Ulm, 2013; Lee and Ellis, 2007; Lumbreras et al., 2010). The AtMKP1 protein is constantly degraded through the proteasome pathway under non-stress conditions, being stabilized *via* phosphorylation upon stress treatment or elicitation with pathogen-associated-molecular-patterns (PAMPs) (Jiang et al., 2017). The posttranslational regulation or turnover of AtMKP2 protein has not been investigated yet. The AtMKP2 has been identified as a positive regulator of oxidative stress tolerance, based on analysis of inducible AtMKP2-RNAi knockdown lines and *mkp2* mutants exhibiting the hypersensitivity to ozone and oxidative agent methyl-viologen, respectively (Lee and Ellis, 2007; Lumbreras et al., 2010). Interestingly, AtMKP2 appeared as the only one among five *Arabidopsis* DSPs that was able to deactivate AtMPK3 and AtMKP6 in response to oxidative stress (Lee and Ellis, 2007). AtMKP2 has also been associated with regulation of immunity–related responses (Lumbreras et al., 2010) and senescence (Li et al., 2012), but the immunity role seems to be dependent on the pathogen life-style. The lack of AtMKP2 correlated with an attenuation of symptoms caused by biotroph *Ralstonia solanacearum*, and accelerated disease progression upon infection with the necrotroph *Botrytis cinerea,* arguing for a distinct role of this phosphatase in regulation of hypersensitive response and cell death (Lumbreras et al., 2010; Vilela et al., 2010).

Oxidative stress is triggered upon plant exposure to both, biotic and abiotic environmental challenges, due to accumulation of reactive oxygen species (ROS). ROS are important players in stress signaling and adaptation responses (Mittler et al., 2022); however, they have to be tightly regulated by ROS scavenging mechanisms to prevent cell damage. Glutathione-ascorbate cycle plays a central role in this process by maintaining the reduced to oxidized glutathione ratio (GSH/GSSG), which is crucial for cellular redox homeostasis (Liu and He, 2017). MAPK and ROS pathways mutually impact each other. On one hand, MAPK cascades can be activated by ROS, on the other hand, MAPK signaling can regulate the expression of ROS–related genes (Boro and Chattopadhyay, 2022; Liu and He, 2017). In *Arabidopsis*, several MAPK pathways can be activated by hydrogen peroxide, including the AtMKK4-AtMPK6, AtMKK3-AtMPK7 and MEKK1-MKK1/2-MPK4 (Doczi et al., 2007; Pitzschke et al., 2009). The AtMPK3 and AtMPK6 can be activated by both, exogenous application of hydrogen peroxide and by intracellular perturbations of ROS upon oxygen deprivation (Chang et al., 2012). Despite obvious interplay of MAPK pathways and ROS signaling, the mechanism of MAPK activation in respect to ROS and oxidative stress remains unclear.

A well described mechanism of sensing ROS is the modification of the redox-sensitive cysteine residues in proteins resulting in sulfenylation, sulfinylation, sulfonylation, or S-glutathionylation (Dalle-Donne et al., 2007; Schieber and Chandel, 2014). While sulfenylation is reversible and may be a part of redox-signaling, the sulfinic and sulfonic modifications are irreversible and cause protein damage. Protein S-glutathionylation is an alternative modification forming a mix disulfide bond between a cysteine residue and glutathione. It plays important role in redox signaling as well as in protecting proteins from irreversible oxidation of protein thiols (Hill and Bhatnagar, 2012). S-glutathionylation is usually considered a modification occurring in response to enhanced production of ROS or oxidation of GSH to GSSG, although it also occurs under normal physiological conditions (Huang and Huang, 2002).

Recently, it has been shown that transgenic plants with increased glutathione content display an oxidative shift in the cytosolic glutathione redox potential and have constitutively active immunity related mitogen-activated protein kinases namely salicylic acid induced protein kinase (SIPK) and wound induced protein kinase (WIPK) (Matern et al., 2015). The mechanism by which glutathione can activate or keep these MAPKs in activated state is not known. In this study, it has been demonstrated that the *A. thaliana* orthologs of WIPK and SIPK, namely AtMPK3 and AtMPK6, exhibit hypersensitivity to the flagellin fragment (flg22) in the *cat2-1* line under conditions causing elevated glutathione and redox perturbations. As AtMPK3 and AtMPK6 are deactivated by AtMKP2 in response to oxidative stress, we hypothesized that AtMKP2 may be affected in the *cat2-1* background. Our results indicate that AtMKP2 remains stable in non-treated *cat2-1* line, but get degraded under severe stress provoked by the treatment of *cat2-1* seedlings with exogenous hydrogen peroxide. The active site cys109 stabilizes the protein under the severe stress conditions and can be glutathionylated *in vitro*.

## Materials and Methods

### Plant material and growth conditions

Plant materials used in this study were *Arabidopsis thaliana* (L.) including wild-type Columbia (Col-0), catalase-deficient mutant *cat2-1* and phytoalexin-deficient mutant *pad2-1*. The seeds were obtained from the Nottingham Arabidopsis Stock Centre (http://nasc.nott.ac.uk). Transgenic mutant lines overexpressing MAPK phosphatase 2 (MKP2) were in the same ecotype backgrounds. Seeds were sterilized, stratificated for 2 days at 4 °C and germinated on ½ Murashige and Skoog (MS) medium including vitamins (Duchefa Biochemie, Haarlem, the Nether-lands) containing 20 g/l sucrose and 8 g/l Microagar (Duchefa Biochemie). Plants were cultivated under white fluorescent light (Lumilux©Cool daylight 865; Osram, Munich, Germany). The photoperiod used in this study is either 8 h light / 16 h night for the short-day condition or 16 h light / 8 hours night for the long-day condition, depending on different experimental setup. The cultivating temperature is 22°C light / 18°C night with the light intensity of 140-200 μmol m^-2^ s^-1^.

### Generation of constructs and transgenic lines

The greengate cloning strategy was used to generate constructs for stable transformation of wild type (wt) and *cat2-1 A. thaliana* lines (Lampropoulos et al., 2013). The greengate modules were kindly provided by Jan Lohmann’s group at the Centre for organismal studies, Heidelberg University. C109S mutation in MKP2 was introduced by the site-directed mutagenesis. The wild-type and mutant versions of MKP2 in constructs were verified by sequencing. For plant transformation, both the wt and the C109S mutant version of MKP2 sequence were fused with c-myc at N-terminus, placed under control of dexamethasone inducible expression system p6xoP/GR_LhG4 (Craft et al., 2005) and introduced into *Agrobacterium tumefaciens* strain C58C1. The primers and constructs are listed in the **Supplementary Table S1 and S2**, respectively.

*Arabidopsis* plants were transformed by dipping the inflorescences into the *Agrobacteria* solution as described by Lampropoulos *et al*. (2013). For plant selection on soil, 0.02 % Basta^TM^ (glufosinate-ammonium; Bayer, Leverkusen, Germany) was used for spraying. For selection on ½ MS plates (with or without sugar) glufosinate-ammonium (Sigma) was used in final concentration of 7.5 µg/ml. T3 seedlings from 100 % Basta resistant T3 seed pools (homozygous for myc-MKP or myc-MKP C109S) were used for experiments on MKP2 protein stability.

### MAPK activation and stress treatments

For MAPK activation experiments, three weeks old *A. thaliana* seedlings, grown on ½ MS plates, were carefully transferred to 24-well-plates (3/well) containing 1 ml of ½ MS liquid medium without sugar and kept overnight in the climate chamber or growth cabinet, under same conditions where plants were before treatments. MAPKs were activated by replacing the MS medium with 100 nM flg22 solution. For MKP2 stability experiments, 2–3 weeks old seedlings grown on 1/2 MS plates were dripped with dexamethasone solution on germination plates to induce the expression of myc-tagged wild-type or mutated version of MKP2. After 6 hours, dexamethasone solution was replaced with cycloheximide, H_2_O_2_ or both together. The treatment solutions were prepared in liquid ½ MS medium as follows: Dexamethasone: 30 µM (from 30 mM stock in ethanol) supplied with 0,005% silwet L-77, cycloheximide: 100 µM (from 10 mM stock in DMSO, final 1% DMSO), H_2_O_2_: 20 mM. For mock treatments, the corresponding solvents were used in the same concentration as for treatments. After treatments, seedling were taken out from wells or plates and the excess of liquid was quickly removed by tissue paper. Thereafter, samples were immediately frozen in liquid N_2_ and kept at -80°C until analyzed.

### Glutathione measurement

Thiols were extracted from 30 mg plant material by phosphate buffer containing either 5 mM DTT for GSH or 5 mM N-ethylmaleimide for GSSG, and 30 mM monobromobimane (dissolved in acetonitrile; Sigma) were added for derivatization as described by Fey *et al*. (2005). Samples were then subjected to Acquity H-class UPLC system with Acquity BEH Shield RP18 column (50 mm x 2.1 mm, 1.7 µm, Waters) and glutathione was quantified according to Liedschulte *et al*. (2010).

### SDS-PAGE and immunoblot analysis of MAPKs and MKPs

Proteins were extracted from N_2_-frozen and ground plant material. For the analysis of MAPK activity and protein levels, the Lacus extraction buffer was used for protein extraction (50 mM Tris-HCl, pH 7.5, 10 mM MgCl2, 15 mM EGTA, 100 mM NaCl, 2 mM dithiothreitol, 1 mM NaF, 1 mM NaMo, 0.5 mM NaVO_3_, 30 mM β-glycerophosphate, and 0.1% Noniondet IGEPAL and 1% anti-protease cocktail (P9599; Sigma)). TEDAS buffer (50 mM Tris-HCl, pH 7.5, 3 mM EDTA, 3 mM dithiothreitol, 10 mM ascorbate and 250 mM sucrose) was used for the analysis of MKP2 stability. Homogenate was centrifuged 2 times at 15,000 x *g* and 4°C for 10 min, and supernatant was taken for protein analysis. If not indicated differently, 15 µg of soluble protein were loaded per lane for immunoblot analysis. For immunodetection of active MAPKs 30 µ of soluble protein were loaded per lane. For the analysis of the whole cell extract, proteins were extracted in 1x reducing ROTI®Load 1 buffer (Carl Roth GmbH + Co. KG, Karlsruhe, Germany) from N_2_-frozen ground tissue powder (5 µL/mg fresh weight), and 10µl of extract were loaded per line for western blots (equivalent to 2 mg fresh weight).

Immunoblot analysis was performed as described by Han *et al*. (2013a). The anti-pTEpY antibody purchased from Cell Signaling (NEB 4370S) was used for detecting phosphorylated MAPK3 and MAPK6 following the manufacturer’s instructions. Total MAPK3, MAPK6 and MKP2 protein were detected with anti-AtMPK3 antibody (Sigma M8318), anti-AtMPK6 antibody (Sigma M7104) and anti-myc antibody (Biolegend 626801), respectively.

### Gene expression analysis

Real-time quantitative (Q) PCR was used for gene expression analysis. Total RNA was extracted from plant materials by a Gene Matrix Universal RNA Purification kit (EurX). First-strand cDNA was synthesized using SuperScript III Reverse Transcriptase (Invitrogen). One reaction was done in total volume of 15 μl including: 1 μl of 5 μM forward and reverse primer solutions, 1.5 μl of 10x Taq Polymerase Buffer, 0.3 μl of 10 mM dNTPs, 0.3 μl Jump start Taq polymerase (Sigma), 0.15 μl SYBR Green (Sigma, dilluted 1:400), 5.75 μl ddH_2_O and 5 μl of cDNA template. Polymerase chain reaction (PCR) was performed following the cycling conditions: denaturation at 95 °C for 6 min, followed by 40 cycles of 95 °C 30 s, 60 °C 20 s, and 72 °C for 30 s. Expression levels of designated genes were normalized to the expression of two reference genes, the clathrin adaptor subunit (AT5g46630) and expressed protein At4g26410 (Czechowski et al., 2005). ΔΔCq method was used for quantification analysis (Livak and Schmittgen, 2001).

### In vitro glutathionylation

Soluble proteins were extracted from leaves of soil-grown 4 weeks old *A. thaliana* plants carrying dexamethasone-inducible myc-tagged wild-type or mutated (C109S) versions of AtMKP2. Transgene expression was induced 12 hours before sampling by spraying the plants with 15 µM dexamethasone in 0.1% ethanol.

For protein extraction, 5 g of tissue were ground at 4°C in 10 ml of extraction buffer containing 0.1 M Tris/HCl, pH 7.5, 1 mM EDTA and 2 mM DTT. Treatment with GSSG-biotin (final conc. 10 µM) and subsequent pull-down of glutathionylated proteins with streptavidin-agarose was performed as described by Dixon *et al*. (2005). The washing steps were conducted with buffer A (20 mM Tris/HCl, pH 6.8, 0.5 M NaCl, 1 mM EDTA) and buffer B (20 mM Tris/HCl, pH 6.8, 6 M urea) to remove nonthiolated proteins binding to the matrix *via* interaction with thiolated proteins. After the final washing step with buffer A, the thiolated proteins were released from beads by adding the loading buffer ROTI®Load 2 (Carl Roth GmbH + Co. KG, Karlsruhe, Germany), supplied with 100 mM DTT. After heating of samples at 95°C for 5 min, equal volumes were loaded on the gel for immunoblot analysis.

For N-ethyl-maleimide (NEM) binding, NEM-biotin was used in final concentration of 30 µM and pull down of proteins interacting with NEM was performed as for GSSG-biotin. As a control, the extract mixture containing the same amounts of wt and mutated MKP2 was treated with nonbiotinylated GSSG and NEM and all steps were repeated in the same way as for GSSG-biotin and NEM-biotin.

### Statistical analysis

Statistically significant differences between *Arabidopsis* lines of wild type and mutant were determined using Student’s t-test in the software package IBM SPSS Statistics. An indicated *P*-value of < 0.05 was regarded as statistically significant.

## Results

### MAPK activity is affected in Arabidopsis line cat2-1 that exhibits a perturbed glutathione homeostasis

Our previous results indicated that perturbed glutathione homeostasis affects regulation of MAPK activity and the signal strength in tobacco (Matern et al., 2015). To prove if this might be a general phenomenon, we examined the activation of defense-related MAP-Kinases in *Arabidopsis thaliana*, AtMPK6 and AtMPK3, in the background of mutant lines *cat2-1* and *pad2-1*, which exhibit conditionally high and permanently low glutathione levels, respectively. The *cat2-1* line has an impaired expression of the class I catalase, crucial for removal of photorespiratory H_2_O_2_ (Queval et al., 2007). This causes a severe oxidative stress when plants grow in ambient air conditions and at irradiance intensity above 100 µmol m^-2^ s^-1^, leading to a plant growth inhibition. Under these conditions *cat2-1* line shows a perturbed intracellular redox homeostasis with high glutathione levels and activated oxidative signaling pathway. On the other side, *pad2-1* mutant is impaired in glutathione synthesis due to mutation in γ-glutamyl cysteine ligase (Parisy et al., 2007).

Interestingly, AtMPK3 and AtMPK6 activation in *A. thaliana* mutant line *cat2-1*, which accumulates high levels of oxidized glutathione was hypersensitive to bacterial flagellin peptide flg22 (**Fig. 1**). The signal for active MAPK3 and MAPK6 was detected after 10 minutes of treatment with flg22 in both Col-0 (wt) and *cat2-1* lines, having a peak after 20 minutes of exposure to flg22. However, the signal was much more abundant in *cat2-1* line and was maintained longer, suggesting a stronger activation of AtMPK3 and AtMPK6 in this genetic background as compared to the control (Col-0). In contrast, the mutant line *pad2-1* that is able to accumulate only 20 % of the glutathione amount usually found in Col-0 background (Parisy et al., 2007), has shown inconsistent results regarding the MAPK activation in response to flg22. The MAPK activation upon flg22 treatment was in most experiments less prominent in *pad2-1* background than in Col-0 (**Fig. 1**), but in some experiments the signal intensity was similar to wild type. This is suggesting a complex role of glutathione in MAPK activation although the *pad2-1* line exhibits enhanced susceptibility to several pathogens indicating an important role of glutathione in defense responses (Parisy et al., 2007). Under the short-day conditions none of the examined lines showed a sustained MAPK activity in the absence of flg22.

**Fig. 1.**
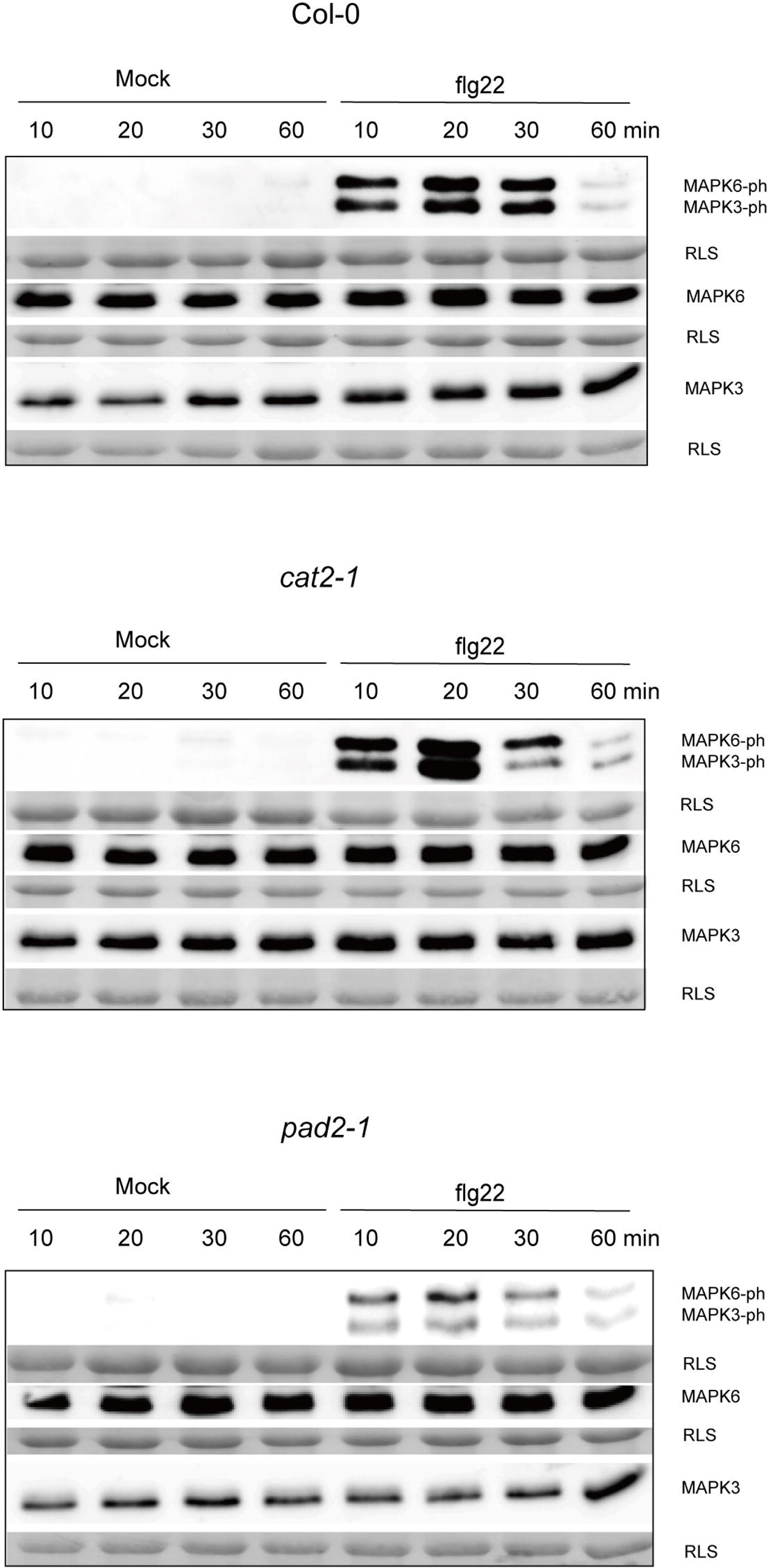
AtMPK3 and AtMKP6 sensitivity to flg22 in Arabidopsis lines Col-0, *cat2-1* and *pad2-1*. Seedlings were treated with 100 nm flg22 for the indicated time intervals. The strongest activation of both, MAPK6 and MAPK3 was detected after 20 minutes Non-treated and mock treatment controls showed no activation of MAPK cascade. Twenty μg of total protein were loaded for detecting MAPK active form and 15 μg for MAPK protein levels. The anti-pTEpY antibody was used for detecting phosphorylated MAPK3 and MAPK6. For detection of the whole protein, the antibodies specific for *Arabidopsis thaliana* MAPK3 and MAPK6 were used. After detection, membranes were stained with amidoblack. Rubisco large subunit (RLS) indicates the loading control. The same results were obtained in four independent experiments.

To answer the question if differences on MAPKs activation pattern rely on the differences in protein level, MAPK protein levels were also analyzed. As shown in **Fig. 2**, MAPK protein levels in *cat2-1* and *pad2-1* are similar to the wild type in both flg22 non-treated and treated samples, indicating that the difference in activity is not caused by differences in protein levels. In addition, gene expression levels of AtMPK3 and AtMPK6 genes as well as genes for MKP1 and MKP2 were not significantly different from the wild type. GSH and GSSG levels as well as the ratio of GSH/GSSG were confirmed in all examined lines (Col-0, *cat2-1* and *pad2-1*) by UPLC analysis (**Supplementary Fig. S1**).

**Fig. 2.**
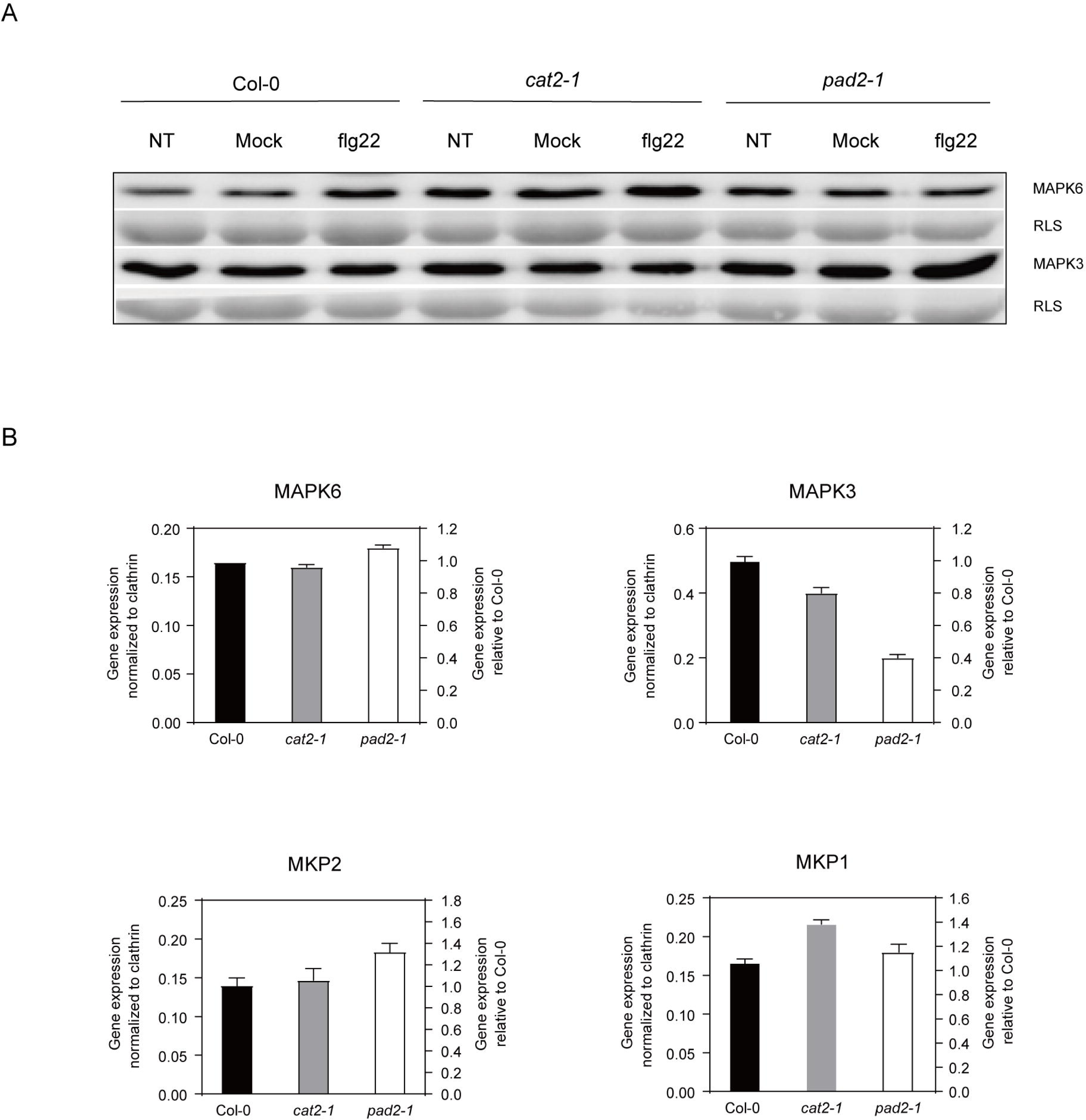
Monitoring MAPK protein and transcript levels, and MKP transcript levels in different Arabidopsis lines. Three weeks old Col-0, *cat2-1* and *pad2-1 Arabidopsis thaliana* seedlings from ½ MS plates were used for the experiments. (A) MAPK3 and MAPK6 activation by mock and flg22 (100 nM) treatments. Twenty μg of total protein was loaded to detect phophorylated MAPK3 and MAPK6 and 15 μg for MAPK3 and MAPK6 protein levels. NT refers to non-treated samples. After detection, membranes were stained with amidoblack. Rubisco large subunit (RLS) indicates the loading control. (B) Relative expression levels of MAPK3, MAPK6, MKP1 and MKP2. Clathrin adaptor subunit and At4g26410 were used as reference genes (Czechowski et al., 2005). Results are represented as mean values of two biological replicates ± standard error.

### MAPK phosphatase 2 stability in cat2-1 line is comparable to that of Col-0

The sustained activation of tobacco WIPK and SIPK in high glutathione lines and hyper-reactivity of the Arabidopsis immunity-related MAPKs in *cat2-1* mutant, prompt us to consider MAPK deactivation as a target of glutathione-dependent regulation. MAPKs can be fully deactivated only by dual-specificity phosphatases that dephosphorylate both, phosphotyrosine and phosphothreonine residues in the MAPK active site. In *Arabidopsis*, only one dual specificity phosphatase MKP2 was shown to be able to deactivate AtMPK6 and AtMPK3 in response to oxidative stress (Lee and Ellis, 2007). MKP2 is the smallest of the *Arabidopsis* MAPK phosphatases (18.4 kD, 167 amino acids) consisting solely of a dual-specificity phosphatase catalytic site and has no other conserved motifs. MKP2 transcript is barely affected by stress treatments (Winter et al., 2007), suggesting a regulation at protein level; however, to our knowledge, there is no information about the posttranslational regulation of MKP2. We hypothesized that the oxidative stress conditions in *cat2-1* background might have destabilized the MKP2 protein, thus changing the magnitude of the MAPK activation in this background.

To examine the protein stability of MKP2, we generated stable homozygous *Arabidopsis* transgenic lines with dexamethasone-inducible expression of the wild-type MKP2 and the mutant version of MKP2 protein (cys109ser) in Col-0 and *cat2-1* backgrounds. The cys109 is the catalytic cysteine residue in the active site of MKP2 (Lee and Ellis, 2007), which is conserved among all PTPs and DSPs and directly involved in MAPK dephosphorylation (Farooq and Zhou, 2004). Due to the low pKa determined to be between 4 - 6 for different PTPs, the catalytic cysteine residues are more susceptible to oxidation than other cysteines (pKa between 8 and 9) (Östman et al., 2011). Moreover, it has been reported that human MPK phosphatase 1 can be inactivated by glutathionylation of the catalytic cys residue and subsequently targeted for ubiquitylation and degradation via proteasome (Kim et al., 2012). In **Fig. 3**, the DSP catalytic site, including the conserved cysteine, is indicated in the alignment of *Arabidopsis* MKP2 protein sequence with those of human MKP1 and *Arabidopsis* MKP1 as examples for DSPs.

**Fig. 3.**
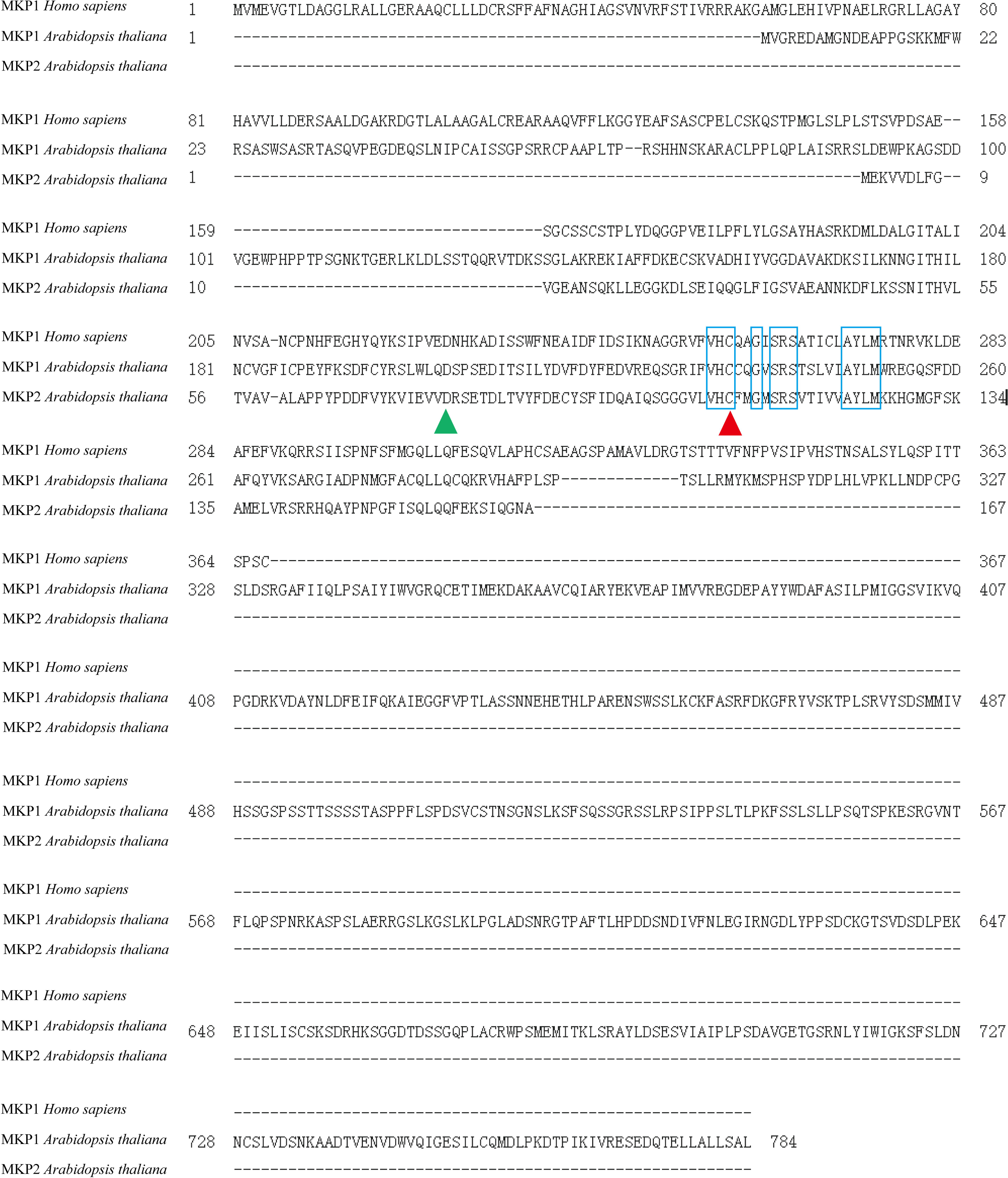
Sequence alignment of human and plant MKPs. Protein sequence of MKP1 from *Homo sapiens* was alligned with MKP1 and MKP2 from *Arabidopsis thaliana* by using Clustal Omega. Conserved cysteine residues in the active site are labeled by the red arrow head. The crucial upstream asp-residue (D78 in AtMKP2) is labeled by the green arrow head. The active site sequence motif VHC-x2-G-x-SRS-x5-AYLM is highlighted by blue frame.

The phenotype of transgenic lines carrying the dexamethasone-inducible, c-myc-tagged MKP2 constructs was similar to untransformed Col-0 and *cat2-1* lines at all developmental stages. Besides, the myc-tagged MKP2 was not detected in the absence of dexamethasone treatment indicating tightly regulated expression of the protein. Upon treatment with dexamethasone, c-myc:MKP2 could be detected in both genetic backgrounds, Col-0 and *cat2-1* (**Supplementary Fig. S2**). The signal started to appear approximately 3 to 6 hours after induction and continued to accumulate for at least 24 hours. The protein abundance differed among the lines, but was not dependent on the genetic background or introduced cys109ser mutation. To examine the stability of MKP2 in the extract, the soluble fraction extracted from dexamethasone induced seedlings was incubated at room temperature for up to 60 minutes without the addition of protease inhibitors. The wild-type version of the protein was detected all over the incubation time, however, the mutated version c-myc:MKP2cys109ser appeared less stable in the extract and was barely detectable after 60 minutes (**Fig. 4**). Altogether the results suggest that in contrast to AtMKP1, AtMKP2 is most likely not rapidly turned-over.

**Fig. 4.**
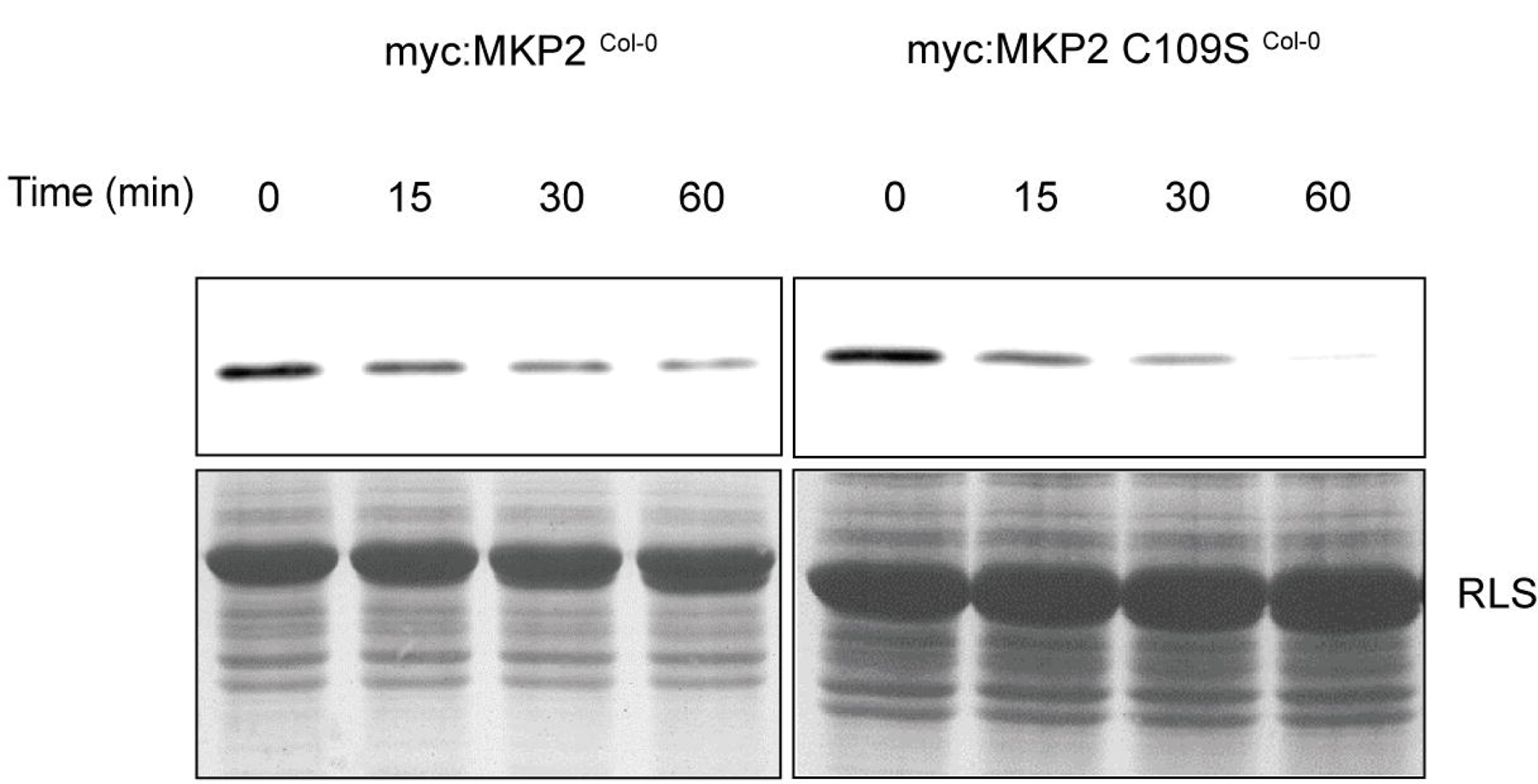
Effect of C109S mutation in MKP2 on protein stability in extract. Soluble proteins were extracted from 18-days-old *Arabidopsis thaliana* seedlings expressing the dexamethasone-induced myc-tagged wild-type, or cys109ser-mutated MKP2 protein. Extracts were divided in four portions and incubated at RT for the indicated time intervals. MPK2 was detected with anti-myc antibody. The equal volumes (corresponding to the 15 µg of total protein as estimated prior to incubation) were loaded for each treatment. Rubisco large subunit (RLS) indicates the loading control. The same results were observed in two independent experiments.

Aiming to increase the stress impact in *cat2-1* line, the light period for seedlings growth was extended from 8 to 16 hours. Under these conditions, *cat2-1* seedlings did exhibit a sustained AtMPK6 activity and, to some extent, sustained AtMPK3 activity (**Fig. 5A**), similar to observations from our previous study on high-glutathione tobacco lines. To examine the stability of c-myc:MKP2 under these conditions, the protein expression in *Arabidopsis* seedlings has been induced by dexamethasone and the turn-over of both wild-type and mutated version of c-myc:MKP2 was monitored in the presence of protein translation inhibitor cycloheximide that was added 6 hours upon dexamethasone induction (**Fig. 5B**). Cycloheximide has disabled the protein accumulation in all genetic backgrounds tested, however, in all examined cases, the protein amount detected after 24 hours incubation with the inhibitor was comparable to the protein amount detected upon 6 hours dexamethasone induction, suggesting that both protein versions, the wild-type and the mutated one, remained stable in both genetic backgrounds (**Fig. 5B**). Although the magnitude of AtMPK3 and AtMPK6 activity seems to be affected in *cat2-1* seedlings, this is most likely not related to the MKP2 protein instability or degradation.

**Fig. 5.**
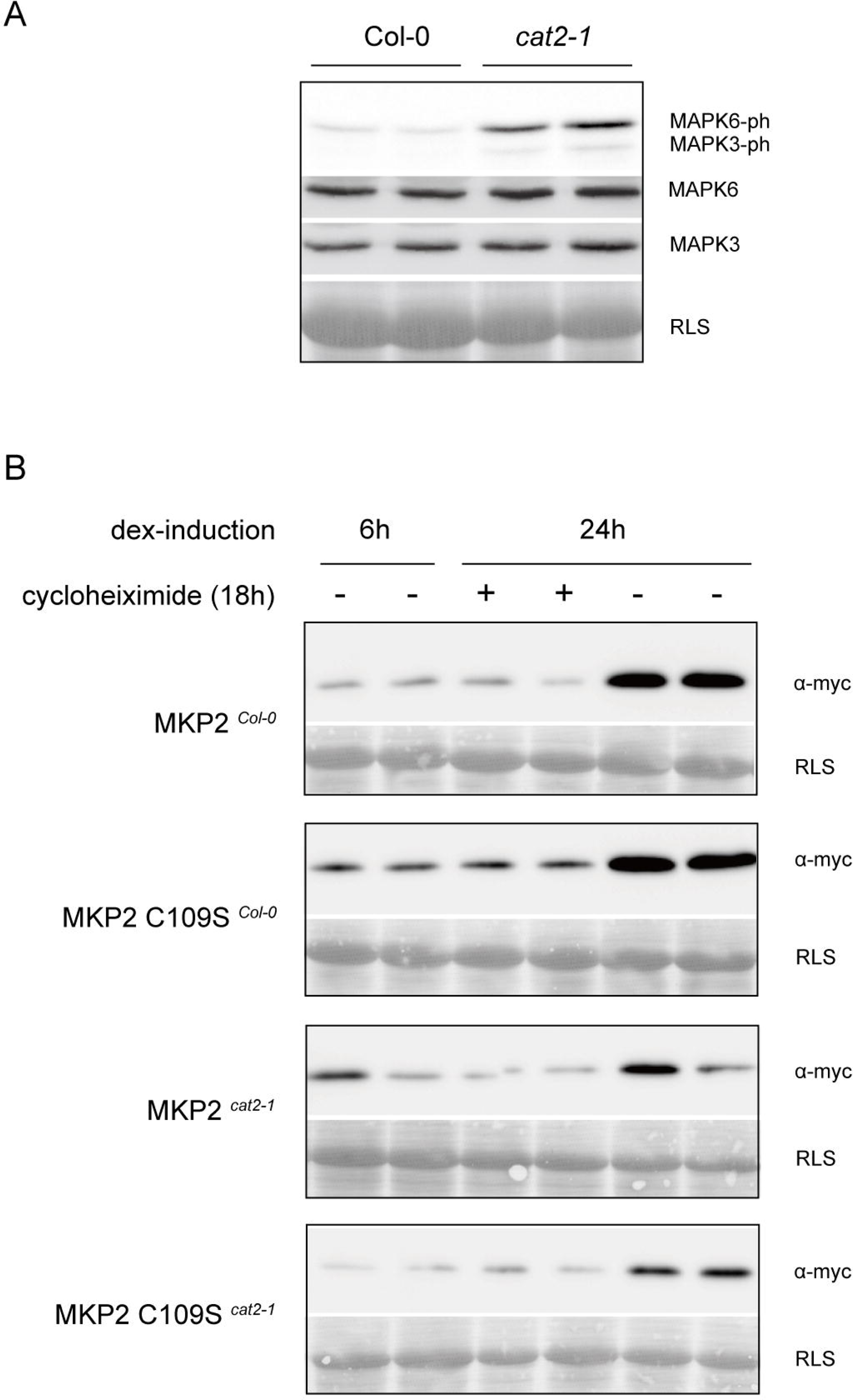
MAPK activity and MKP2 stability in Col-0 and *cat2-1* under long photoperiods. (A) Steady-state MAPK activity in non-induced and non-treated Col-0 and *cat2-1* seedlings. Immunodetection of the phosphorylated MAPK3/6, AtMPK3 and AtMPK6 was performed as described for Fig.1. (B) Cycloheximide treatment for determination of MKP2 stability in different genetic backgrounds. 14 days old seedlings germinated on ½ MS plates were dripped with dexamethasone (30µM) solution to induce the expression of myc-tagged wild-type or mutated MKP2. After 6 hours, dexamethasone solution was replaced with cycloheximide (100 μmol) or mock solution and seedlings were treated for another 18 hours. Whole-cell extracts from treated samples were subjected to immunoblot analysis with anti-myc antibody (sample volume equivalent to 2 mg of fresh weight was loaded per lane). Rubisco large subunit (RLS) indicates the loading control (amidoblack membrane staining). The same results were obtained in five independent experiments.

### MAPK phosphatase 2 is unstable under severe oxidative stress

In an effort to examine the effect of exogenous oxidative stress on MKP2 stability, H_2_O_2_ treatment was applied to seedlings growing under long-day conditions, and the c-myc:MKP2 turn-over was monitored in the Col-0 and *cat2-1* backgrounds (**Fig. 6**). After 6 hours of dexamethasone-driven protein expression, protein translation was inhibited *via* cycloheximide and seedlings were treated for 18 hours with H_2_O_2_. In the wild type, c-myc:MKP2 remained stable, but severe stress in the *cat2-1* caused degradation of c-myc:MKP2 in this background (**Fig. 6B**). Furthermore, we examined if change of the active site cys109 for serine would prevent the MKP2 degradation under severe stress, as this cysteine residue could be a target of protein oxidation. For this, protein stability of both the wild-type and the mutated version of c-myc:MKP2 has been investigated in *cat2-1* background upon H_2_O_2_ treatment, while monitoring at the same time a distribution of the protein between the soluble and microsomal fraction of the extract (**Fig. 6C**). H_2_O_2_ *per se* did not prevent the dexamethasone-induced-accumulation of MKP2 in the *cat2-1*, however, the treatment with the protein translation inhibitor cycloheximide and H_2_O_2_ at the same time caused a decrease in the wild-type version of the protein in both soluble and microsomal fraction. Surprisingly, the absence of the catalytic cys109 in the mutated version of the protein correlated with more prominent protein degradation, which was apparent in the soluble and even more in the microsomal fraction of the extract. In contrast to our initial hypothesis, this was suggesting a rather stabilizing role for the cys109 residue under the conditions of severe stress. Moreover, as MKP2 is a soluble protein, the association with microsomal fraction could have happened due to interaction with other molecules, which might be disrupted by replacing cys109 with serine residue.

**Fig. 6.**
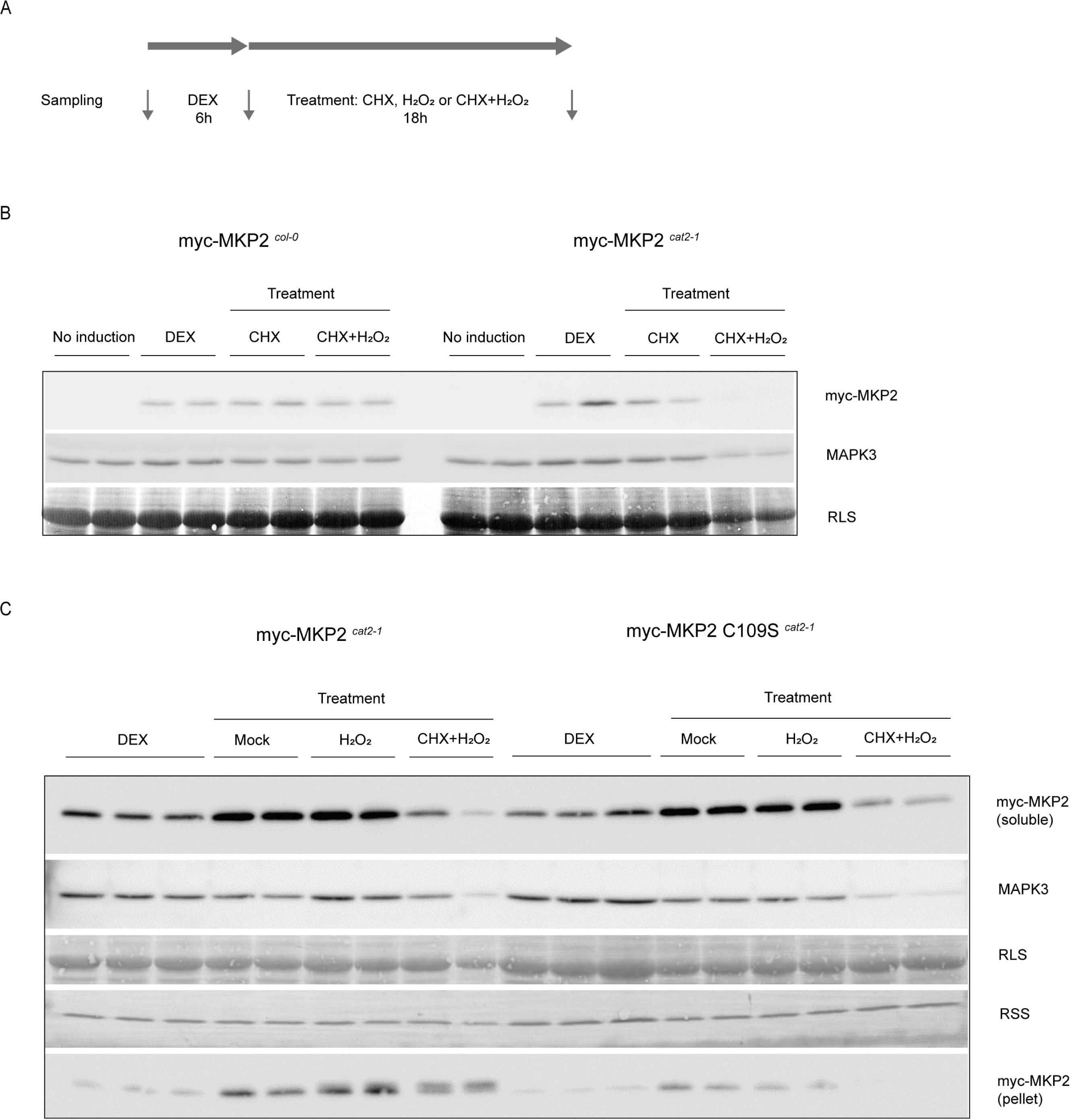
Effect of H_2_O_2_ on MKP2 stability in Col-0 and *cat2-1* lines. (A) The experimental setup. 17-days-old seedlings grown on ½ MS plates in long-day conditions were treated as described for Fig.5. (B) Stability of the myc-tagged wild-type MKP2 in Col-0 and *cat2-1* backgrounds upon treatment with cycloheximide and H_2_O_2_. Whole-cell extracts from treated samples were subjected to immunoblot analysis with anti-myc antibody, where equal sample volumes were loaded for each sample, equivalent to 2 mg of fresh weight per lane. (C) Stability of the wild-type and cys109ser-mutated MKP2 in *cat2-1* background. Upon protein extraction, soluble and microsomal protein fractions were analyzed separately. Pellets were resuspended in the same volumes of extraction buffer, as used for extraction (1.4 µl/mg fresh weight). The equal volumes of denatured soluble and microsomal fractions were loaded per lane for each sample, corresponding to the 15 µg of protein in the soluble fraction. MAPK3 protein amount was also monitored. Rubisco large subunit (RLS) and rubisco small subunit (RSS) indicate the loading control.

As mentioned above, the catalytic cysteine residues in PTPs and DSPs are more susceptible to oxidation than other cysteines, which may affect the protein activity and stability (Östman et al., 2011). However, the reversible S-glutathionylation, which targets cysteines in the basic environment as well, might protect proteins by preventing from sulfhydryl overoxidation or proteolysis (Grek et al., 2013). To investigate if MKP2 can be modified by glutathione, S-glutathionylation experiment was conducted for MKP2 *in vitro* (**Fig. 7**). Wild-type MKP2 protein and the MKP2cys109ser mutant were treated with GSSG-biotin and incubated with streptavidin agarose to pull down glutathionylated proteins by affinity purification. The wtMKP2, but not the MKP2cys109ser, remained bound to the streptavidin upon binding to GSSG-biotin (**Fig. 7**, upper panel). In the N-ethyl-maleimide (NEM) binding experiment, both wild-type and cys109ser-mutated MKP2 were able to bind NEM and were pulled down with streptavidin agarose (**Fig. 7**, middle panel). Since the cys109ser-mutated MKP2 protein still has an intact cysteine at position 91, the protein was able to bind NEM but apparently could not be glutathionylated *in vitro*. Altogether, our results indicate that the catalytic cysteine residue (cys109) in the active site of MKP2 is important for the protein stability under severe stress and may be a target for S-glutathionylation.

**Fig. 7.**
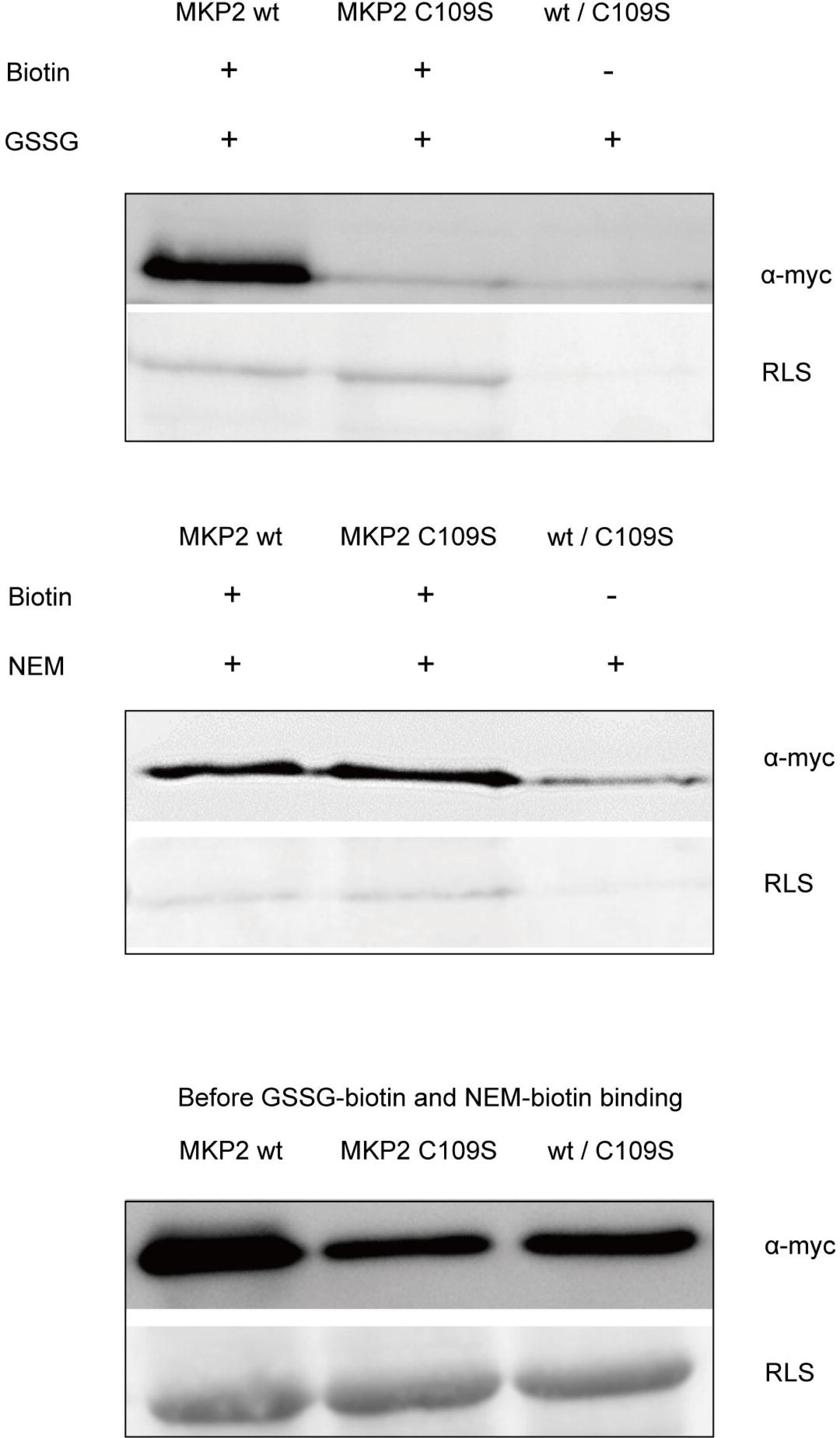
Identification of glutathionylated MKP2. Soluble proteins were extracted from leaves of soil-grown 4 weeks old *Arabidopsis thaliana* plants expressing myc-tagged wild-type (MKP2 wt) or mutated (MKP2 C109S) versions of AtMKP2 (5g fresh weight / 10 ml extraction buffer as described in Materials and Methods). Protein extracts were treated with biotin-labeled GSSG (upper panel) or biotine-labeled NEM (middle panel) and biotinylated proteins were pulled down with streptavidin beads. As a control, the wild-type and mutated versions of the MKP2 protein were mixed and incubated with nonbiotinylated GSSG and NEM to exclude the nonspecific binding to streptavidin (wt/C109S lane). Upon affinity purification, the bound proteins were eluted from strepatvidin beads and equal volumes of eluted protein samples were analyzed by immunoblot with anti-myc antibody (α-myc). Rubisco large subunit (RLS) indicates the loading control. The immunoblot of aliquots from protein extracts prior to binding GSSG-biotin or NEM-biotin is shown on the lower panel (20 µg of total protein were loaded per lane).

## Discussion

### MAPK regulation in cat2-1 background

Glutathione is an indispensable metabolite with diverse functions in living organisms including plants. Among these manifold roles, it has been demonstrated that it takes part in responses to pathogens (Noctor et al., 2012). Due to its very complex nature and involvement in various physiological processes, the role of glutathione in plant immunity is still not fully understood. The earlier study has shown that transgenic tobacco plants with enhanced glutathione biosynthesis and an oxidative shift in the cytosolic glutathione redox potential display a sustained activation of immunity-related MAPKs, along with an overall moderately elevated defense response (Matern et al., 2015). Here we demonstrated that the activity of *Arabidopsis* PAMP-responsive MAPKs, AtMPK3 and AtMPK6 were affected in a photorespiratory mutant *cat2-1*, which exhibited elevated glutathione levels, perturbed glutathione homeostasis and activation of oxidative stress pathways, when grown in ambient air. Both MAPKs were hypersensitive to twenty-two amino acid flagellin fragment (flg22) in *cat2-1* seedlings under ambient conditions and 8 h photoperiods (short-day), whereas under 16 h photoperiods (long-day), a faint sustained activity of both MAPKs could be observed even without flg22 treatment. Interestingly, the perturbation of stress responses in *cat2-1* mutant grown in ambient air is additionally affected by photoperiods (Queval et al., 2007). High glutathione levels trigger, under long-day photoperiods but not under short-day ones, lesions on leaves, a comprehensive activation of salicylic acid- and jasmonate-dependent defense responses, and an increase in defense metabolites camalexin and scopoletin (Chaouch et al., 2010; Han et al., 2013b). Moreover, *cat2-1* seedlings exert an augmented callose deposition upon PAMP treatment under these conditions (Luna et al., 2011). The synthesis of camalexin, the major phytoalexin in *Arabidopsis*, and the deposition of callose, crucial for cell wall reinforcement upon pathogen attack, are both regulated by the AtMPK3/AtMPK6 cascade. The activation of AtMPK3/AtMPK6 is sufficient to induce camalexin synthesis in the absence of pathogen attack and it is compromised in *mpk3* and *mpk6* mutants (Ren et al., 2008). On the other side, constitutive activation of AtMPK3 and AtMPK6 by upstream kinase MEK5 is sufficient for callose deposition, whereas the *mpk3* mutant and to a smaller extent *mpk6* mutant show reduced flg22-induced callose accumulation (Frei dit Frey et al., 2014; Zhang et al., 2007). Taken together, this suggests that the defence-related responses reported previously for *cat2-1* line could at least partially be caused by the sustained AtMPK3 and AtMPK6 activity that we observed under long-day conditions. In contrast, under the short-day photoperiods, both MAPKs were not continuously active, which corresponds to the absence of a sustained defence response under these conditions as observed by Queval *et al*. (2007); however, they were hypersensitive to flg22, suggesting that under short photoperiods the defences might be primed. This hypothesis is supported by the earlier study of Pastor *et al*. (2013), who observed an enhanced accumulation of pathogen induced camalexin in *cat2-1* line as compared to the wild type after infection with fungus *Plectosphaerella cucumerina* in short-day conditions. Intriguingly, short and long photoperiods impact glutathione homeostasis in *cat2-1* differently (Queval et al., 2007). Although the ambient-air-grown *cat2-1* line exhibits higher glutathione levels and more oxidized glutathione pool than wild type in both conditions, under short photoperiods (8 h) these parameters are more pronounced than under long-day conditions. It is conceivable that differences in glutathione levels and oxidative stress levels might at least partially be the reason for differences in defence responses in *cat2-1* under both conditions.

In mammalian cells and yeasts, MAPK pathways are redox-regulated at different levels through direct cysteine oxidation events (Latimer and Veal, 2016). The human MAPKK kinase ASK1, which is upstream of both p38 and JNK MAPKs in response to a variety of stimuli undergoes redox control *via* interaction with thioredoxins and peroxiredoxins (Jarvis et al., 2012). Although in plant MAPK modules such a regulation mechanism has not yet been reported, it was shown that the oxidative signal inducible kinase (OXI1) is required for activation of AtMPK3 and AtMPK6 (Rentel et al., 2004), whereas several MAPKKKs in plants can be activated *via* H_2_O_2_, however, it remains to be elucidated, if any of them can act as redox sensor (Liu and He, 2017). Interestingly, three Arabidopsis MAPKs emerged in a screen for the substrates of H_2_O_2_-dependent protein sulfenylation, including the H_2_O_2_-responsive AtMPK4 and AtMPK7, as well as AtMPK2 (Waszczak et al., 2014); but, the biological significance of this modification is still not known.

The enhanced amplitude of AtMPK3 and AtMPK6 activities in *cat2-1* background as well as their prolonged activity upon flg22 stimulation prompt us to consider plant MAPK phosphatases as potential redox-sensitive targets. Due to their redox-sensitive catalytic cysteine residue, MAPK phosphatases may become oxidized by H_2_O_2_ generated in response to a variety of stimuli (Östman et al., 2011), leading to their reversible or irreversible inactivation and subsequent degradation.

### Posttranslational regulation of MAPK-phosphatase 2 (MPK2) under oxidative stress conditions and possible implications for MAPK regulation

MAPK phosphatases play important roles in the regulation of many physiological responses with respect to MAPK cascades. Relatively low number of plant MAPK phosphatases as compared to the number of MAPK modules in plants suggests that a particular MKP will be engaged in different MAPK cascades (Bartels et al., 2010). Consequently, the molecular characterization of MAPK phosphatases may facilitate understanding the function of MAPK cascades. The activity of mammalian and yeast MAPK phosphatases, which are all dual specificity phosphatases, has been comprehensively studied and indicated the regulation by transcriptional induction, protein stability and catalytic activation (Liu et al., 2007). The protein stability is mainly affected by phosphorylation events, although glutathyionylation with subsequent degradation over proteasome pathway has been reported as well (Kim et al., 2012; Liu et al., 2007).

In *A. thaliana*, only two MAPK-related dual specificity phosphatases, MKP1 and MKP2, are able to fully deactivate AtMPK3 and AtMPK6, however, only MKP2 is able to deactivate AtMPK3 and AtMPK6 in response to oxidative stress (Lee and Ellis, 2007). The MKP2 transcript levels do not seem to be under a substantial differential regulation due to stress treatments (Winter et al., 2007), and have similar extent of expression in wild type and *cat2-1* line (Fig. 2). Therefore, we were monitoring the MKP2 protein turn-over in *cat2-1* background, under the conditions which provoked hypersensitivity or sustainable activity of the AtMPK3 and AtMPK6. For this, we expressed the dexamethasone-inducible wild-type MKP2 as well as a mutated version of the protein, having the catalytic cysteine replaced for serine, in both wild type and *cat2-1*. Our results indicated that both protein forms were stable in both genetic backgrounds tested, under short and long photoperiods, indicating MKP2 is a rather stable protein under the conditions of oxidative stress. Only under severe oxidative stress, provoked by the exogenous H_2_O_2_ treatment of the *cat2-1* line, which is particularly sensitive to the treatment due to deprivation of catalase 2, the MKP2 protein was partially degraded. This is different to MKP1, and several human MAPK phosphatases which are constantly turned-over and stabilized *via* phosphorylation (Jiang et al., 2017). Interestingly, it has been shown previously that the activity of both, MKP1 and MKP2 is specifically increased in the presence of their biological substrates, AtMPK6 for AtMKP1 and AtMPK3 or AtMPK6 for AtMKP2 (Lee and Ellis, 2007; Park et al., 2011). Thus, it is likely, that the plant MKP2 protein rather undergoes the regulation at the level of enzyme activity, instead of being regulated at transcript or protein level.

Sulfenylation of the catalytic nucleophilic cysteine leads to the inhibition of protein tyrosine phosphatases (PTPs), including the dual-specificity phosphatase subfamily of PTPs in mammals and yeasts, and represent, in addition to phosphorylation, another important mode of regulation for MAPK phosphatases (Conte and Carroll, 2013). The highly conserved catalytic domain suggests, that such modifications may take place in plant proteins as well, but they have not been described for the subfamily of plant dual specificity MAPK phosphatases yet (Bheri et al., 2021). However, *Arabidopsis thaliana* protein tyrosine phosphatase 1 (AtPTP1), the only Tyr-specific PTPase present in this plant was shown to be directly inhibited by H_2_O_2_ and can be protected by S-nitrosation on its catalytic cysteine (Nicolas-Francès et al., 2022). Even though MKP2 seems to be a rather long-lived protein, our experiments show, that the severe stress does impact MKP2 stability, and replacement of catalytic cysteine with serine residue causes even more pronounced degradation. This indicates that the catalytic cysteine could somehow stabilizes the protein, either relating to protein-protein interactions or *via* modifications. We could demonstrate that MKP2 can be glutathionylated on cys109 *in vitro*, but further experiments are necessary to elucidate the regulation of this MAPK phosphatase *in vivo*. Although the cys109 is crucial for the activity, and any modification on that cys would cause inactivation of phosphatase, the reversible S-glutathionylation could protect the protein against overoxidation (Grek et al., 2013). Moreover, this could explain the hyperactivity and sustainable activity of MAPKs under the oxidative stress conditions. On the other side, the studies on biological function of MKP2 suggest a role in a negative regulation of hypersensitive response, cell death and senescence (Li et al., 2012; Lumbreras et al., 2010). In all of these processes, ROS play a crucial role and demand oxidative stress-resilient regulators, which ensure that physiological transitions proceed in a controlled manner. Thus, further studies are necessary to understand the role of protein modifications in these signaling networks.

## Conclusions

In the current study, we investigated the mechanisms of MAPK activation in the background of elevated glutathione levels and a more oxidized glutathione pool. The focus was on MKP2 regulation, as it dephosphorylates MAPK and exerts redox sensitivity. The results indicate hypersensitivity of MPK3 and MPK6 to the immunity defense inducer flg22 in Arabidopsis mutant line *cat2,* which accumulates high levels of oxidized glutathione. This activation pattern does not rely on MAPK transcriptional or protein abundance, and it is not related to the instability or degradation of MKP2. MKP2 stability in the *cat2* line is comparable to that of Col-0 in seedlings. Further results show that MKP2 can be modified by glutathionylation on cys109 in the active site. The findings of this study are crucial in understanding MAPK-MKP regulation mechanisms under altered oxidation states.

## Acknowledgements

The authors are grateful to Prof. Jan Lohmann and his group at the Centre for Organismal Studies at Heidelberg University for sharing their expertise and clones for the Greengate cloning strategy, and to colleagues at the Metabolomics Core Technology Platform (MCTP) of the Heidelberg University for technical support. The authors would like to thank Rebecca Vazquez for her support in repeating the experiments on MAPK activity and transcript levels.

## Author contributions

TPB and TR: conceptualization; YY, SM, HS, MHV and TPB: performing the experiments; YY and TPB: writing the manuscript; all authors edited and reviewed the manuscript.

## Data availability

All data supporting the findings of this study can be found within the paper and its supplementary data published online.

